# A self-administered, artificial intelligence (AI) platform for cognitive assessment in multiple sclerosis (MS)

**DOI:** 10.1101/611335

**Authors:** Seyed-Mahdi Khaligh-Razavi, Maryam Sadeghi, Mahdiyeh Khanbagi, Chris Kalafatis, Seyed Massood Nabavi

**Affiliations:** Cognetivity ltd, London, UK; Department of Brain and Cognitive Sciences, Cell Science Research Center, Royan Institute for Stem Cell Biology and Technology, ACECR, Tehran, Iran; University of Tehran, Tehran, Iran; South London & Maudsley NHS Foundation Trust, London, UK; Department of Old Age Psychiatry, Kings College London, London, UK

**Keywords:** Multiple sclerosis, BICAMS, digital biomarkers, Integrated Cognitive Assessment (ICA), language-independent, Artificial Intelligence (AI)

## Abstract

**Background:** Cognitive impairment is common in patients with MS. Accurate and repeatable measures of cognition have the potential to be used as a marker of disease activity. We developed a 5-minute computerized test to measure cognitive dysfunction in patients with MS. The proposed test –named Integrated Cognitive Assessment (ICA)– is self-administered and language-independent.

**Objective:** To determine ICA’s validity as a digital biomarker for assessing cognitive performance in MS.

**Methods:** 91 MS patients and 83 healthy controls (HC) took part in substudy 1, in which each participant took the ICA test and the Brief International Cognitive Assessment for MS (BICAMS). We assessed ICA’s test-retest reliability, its correlation with BICAMS, its sensitivity to discriminate patients with MS from the HC group, and its accuracy in detecting cognitive dysfunction. In substudy 2, we recruited 48 MS patients, and examined the association between the level of serum neurofilament light (NfL) in these patients and their ICA scores.

**Results:** ICA demonstrated excellent test-retest reliability (r=0.94), with no learning bias (i.e. no significant practice effect); and had high level of convergent validity with BICAMS. ICA was sensitive in discriminating the MS patients from the HC group, and demonstrated a high accuracy (AUC = 95%) in discriminating cognitively normal from cognitively impaired participants. Additionally, we found a strong association (r=-0.79) between ICA score and the level of NfL in MS patients.

**Conclusions:** ICA can be used as a digital biomarker for assessment and monitoring of cognitive performance in MS patients. In comparison to standard cognitive tools for MS (e.g. BICAMS), ICA is shorter in duration, does not show a learning bias, is independent of language, and takes advantage of artificial intelligence (AI) to identify cognitive status of patients more accurately. Being a digital test, it further has the potential for easier electronic health record or research database integration.

## 1. Introduction

Multiple sclerosis (MS) can cause demyelination and neurodegeneration in patients ^1^. Therefore, cognitive dysfunction is common in MS patients (40-70% of these patients are reported to have cognitive impairment ^2^), and is associated with a higher risk of disease progression in the subsequent years ^2^. Cognitive impairment can have significant negative impacts on several domains of activities of daily living, such as social functioning, employment ^3^ and driving ^4^.

The Brief International Cognitive Assessment for MS (BICAMS)^5,6^ is a cognitive assessment battery for detecting cognitive dysfunction in MS patients. The BICAMS battery includes tests of mental processing speed and memory, and takes about 15 to 20 minutes to administer and score. We used BICAMS in this study as a standard reference test to measure the efficacy of the proposed ICA test in detecting cognitive impairment in MS patients.

Cognition has the potential to be used as a marker of disease progression or treatment efficacy in MS ^7^. When patients report a cognitive problem, they are describing a change in function from a previous level; however, the majority of cognitive tests, due to a learning bias ^8,9^, cannot be used for frequent monitoring of cognitive performance. On the other hand, neuroimaging and fluid biomarkers of disease activity ^10–12^, while more accurate, are less suitable for frequent monitoring of disease progression, and more difficult to integrate into routine clinical practice. In this study, we propose an AI-assisted digital biomarker of cognitive function, appropriate for monitoring the disease activity.

It is documented that the afferent visual system is highly vulnerable to MS^13^. Furthermore, deficit in information processing speed (IPS) is the most prevalent cognitive impairment in MS, and can affect speed of sensory, motor and cognitive processes ^14^. We therefore designed an iPad-based rapid visual categorization task ^15–17^, called the Integrated Cognitive Assessment (ICA), that primarily assesses IPS in visuo-motor domains. The task is designed to give a sensitive, repeatable measure of IPS, and is additionally shown to be correlated with other cognitive domains, such as visual memory and visuospatial ^18^. The test is software-based, self-administered and is shown to have little dependency on participant’s language, and is not confounded by participants’ different level of education ^18^.

In this study, we investigate ICA’s validity as a digital biomarker for assessing cognitive performance in MS. We report results for convergent validity between BICAMS and ICA, test-retest reliability, ICA correlation with serum NfL, effect of repeated exposure to the tests (i.e. learning effect), sensitivity to detecting cognitive impairment, and the accuracy of the ICA test in discriminating MS patients from healthy controls (HC).

## 2. Methods

### 2.1 ICA test description and the scientific rationale behind the test

The ICA test is a rapid visual categorization task with backward masking ^15,16,19^. The test takes advantage of the human brain’s strong reaction to animal stimuli ^20,21^. One hundred natural images (50 animal and 50 non-animal) are carefully selected, with various levels of difficulty, and are presented to the participants in rapid succession. Images are presented at the center of the screen at 7° visual angle. In some images the head or body of the animal is clearly visible to the participants, which makes it easier to detect. In other images the animals are further away or otherwise presented in cluttered environments, making them more difficult to detect. Few sample images are shown in Figure 1. We used grayscale images to remove the possibility of some typical color blindness affecting participants’ results. Furthermore, color images can facilitate animal detection solely based on color, without fully processing the shape of the stimulus. This could have made the task easier and less suitable for detecting less severe cognitive dysfunctions.

**Figure 1.**
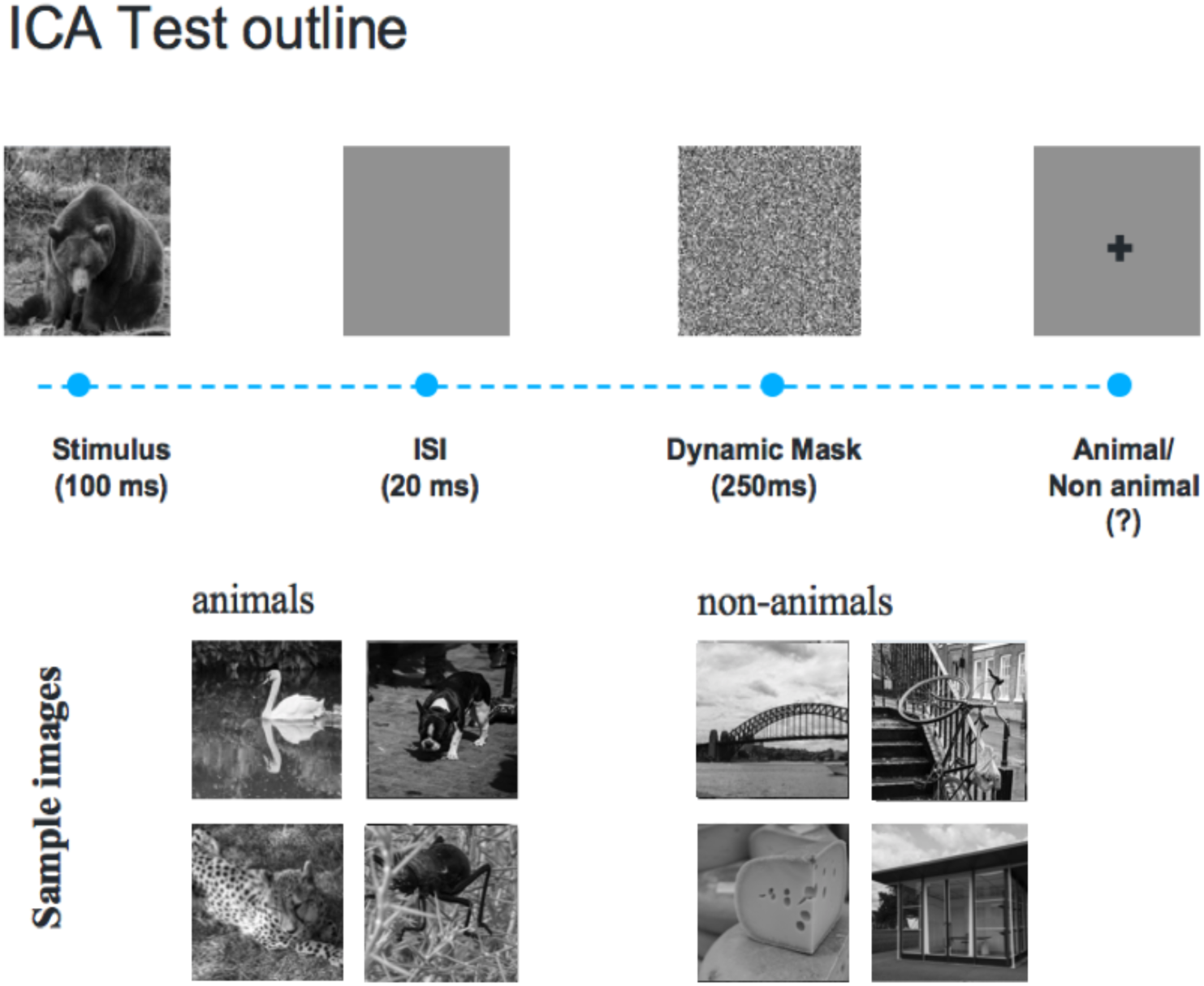
The ICA test pipeline. One hundred natural images (50 animal and 50 non-animal) with various levels of difficulty are presented to the participants. Each image is presented for 100 ms followed by 20 ms inter-stimulus interval (ISI), followed by a dynamic noisy mask (for 250 ms), followed by subject’s categorization into animal vs. non-animal. Few sample images are shown for demonstration purposes.

The strongest categorical division represented in the human higher level visual cortex appears to be that between animates and inanimates ^22,23^. Studies also show that on average it takes about 100ms to 120ms for the human brain to differentiate animate from inanimate stimuli ^21,24,25^. Following this rationale, each image is presented for 100 ms followed by a 20 millisecond inter-stimulus interval (ISI), followed by a dynamic noisy mask (for 250 ms), followed by subject’s categorization into animal vs non-animal (Figure 1). Shorter periods of ISI can make the animal detection task more difficult and longer periods reduce the potential use for testing purposes as it may not allow for the detection of less severe cognitive impairments. The dynamic mask is used to remove (or at least reduce) the effect of recurrent processes in the brain ^26,27^. This makes the task more challenging by reducing the ongoing recurrent neural activity that could artificially boost subject’s performance; it further reduces the chances of learning the stimuli. For more information about rapid visual categorization tasks refer to Mirzaei et al., (2013) ^16^.

The ICA test starts with a different set of 10 test images (5 animal, 5 non-animal) to familiarize participants with the task. These images are later removed from further analysis. If participants perform above chance (>50%) on these 10 images, they will continue to the main task. If they perform at chance level (or below), the test instructions will be presented again, and a new set of 10 introductory images will follow. If they perform above chance in this second attempt, they will progress to the main task. If they perform below chance for the second time the test is aborted.

#### Backward masking

To construct the dynamic mask, following the procedure in (Bacon-Macé and colleagues, 2005) ^15,16^, a white noise image was filtered at four different spatial scales, and the resulting images were thresholded to generate high contrast binary patterns. For each spatial scale, four new images were generated by rotating and mirroring the original image. This leaves us with a pool of 16 images. The noisy mask used in the ICA test was a sequence of 8 images, chosen randomly from the pool, with each of the spatial scales to appear twice in the dynamic mask.

### 2.2 Brief International Cognitive Assessment for MS (BICAMS)

The BICAMS battery consists of three standard pen-and-paper tests, measuring speed of information processing, visuo-spatial memory and verbal learning.

#### Symbol Digit Modalities Test (SDMT)

The SDMT is designed to assess speed of information processing, and takes about 5 minutes to administer ^28^.

#### California Verbal Learning Test -2^nd^ edition (CVLT-II)

The CVLT-II test ^29,30^ begins with the examiner reading a list of 16 words. Participants listen to the list and then report as many of the items as they can recall. Five learning trials of the CVLT-II are used in BICAMS^5^, which takes about 10 minutes to administer.

#### Brief Visual Memory Test–Revised (BVMT-R)

The BVMT-R test assesses visuo-spatial memory ^31,32^. In this test, six abstract shapes are presented to the participant for 10 seconds. The display is removed from view and patients are asked to draw the stimuli via pencil on paper manual responses. The test takes about 5 minutes to administer.

### 2.3 Participants

In total, 174 volunteers took part in substudy1 (Table 1): 91 patients diagnosed with multiple sclerosis (MS), and 83 age, gender and education matched healthy controls. 48 MS patients took part in substudy2 (Table 2). Of all participants 25 attended both substudies. Participants’ age varied between 18 and 65. The study was conducted according to the Declaration of Helsinki and approved by the local ethics committee at Royan Institute. Informed consent was obtained from all participants. Patient participants were consecutively recruited from the outpatient clinic of the Aria Medical Complex for MS in Tehran, Iran. Patients were diagnosed by a consultant neurologist according to the McDonald diagnostic criteria (2010 revision)^33^. Healthy controls (HC) were recruited through local advertisement.

**Table 1.** Demographic and disease related information for participants in substudy 1.

| Characteristic | MS (*n*=91) | HC (*n*=83) | *p*-value |
| --- | --- | --- | --- |
| Age –mean years ±SD | 37.24 ±10.2 | 36 ±10 | 0.42 |
| Education in years –mean ±SD | 14.21 ±3.16 | 14.81 ±2.5 | 0.16 |
| Gender (%female) | 75 (82%) | 58 (70%) | 0.052 |
| Disease Duration (in years) | 6.8 |  |  |
| Disease course |  |  |  |
| Relapsing remitting | 83 (91%) |  |  |
| Secondary progressive | 6 (7%) |  |  |
| Primary progressive | 2 (2%) |  |  |
| EDSS –mean ±SD | 1.27 ±1.8 |  |  |

P-values come from a two-sample t-test. MS: Multiple Sclerosis. HC: Healthy Controls. SD: standard deviation.

**Table 2.**
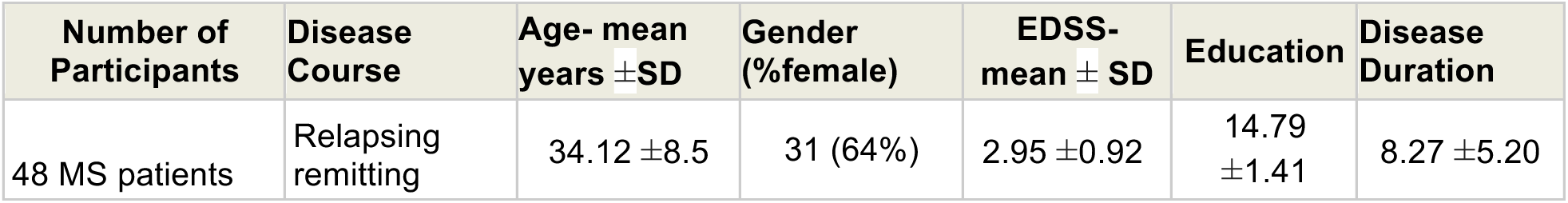
Demographic and disease related information for participants in substudy 2.

Participants’ exclusion criteria included: Severe depression and other major psychiatric comorbidities, presence of neurological disorders and medical illness that independently affect brain function and cognition (other than MS for the patient group), visual problems that cannot be corrected with eye-glasses such that the problem prevents participant from reading, upper limb motor dysfunction, history of epileptic seizures, history of illicit substance and/or alcohol dependence.

For each participant, clinical characteristics of MS subtype, information on age, education and gender were also collected. We quantified participant disability and disability progression over time by utilising the Expanded Disability Status Scale (EDSS).

### 2.4 Study procedures

#### Substudy 1

174 participants (Table 1) took the iPad-based ICA test and the pen-and-paper BICAMS test, administered in random order. The same researchers who administered the BICAMS, directed participants on how to take the iPad ICA test. In this substudy we investigated convergent validity of the ICA test with BICAMS, ICA’s test-retest reliability and the sensitivity and specificity of the ICA platform in detecting cognitive impairment in MS.

To measure test-retest reliability for the ICA test, a subset of 21 MS and 22 HC participants were called back after five weeks (± 15 days) to take the ICA test as well as the SDMT. The subset’s characteristics were similar to the primary set in terms of age, education and gender ratio. For both SDMT, and ICA, the same forms of the tests were used in the re-test session. Note that in the ICA test, while the images were the same, their presentation order randomly changes in every administration.

#### Substudy 2

In this substudy, we investigated ICA’s correlation with the level of serum NfL in 48 MS patients (Table 2). Participants took the iPad-based ICA test and the pen-and-paper SDMT test, administered in random order. ICA and SDMT were administered in the same session, but blood samples were collected in another visit with a gap of 2-3 days in between.

Blood samples were collected in tube for serum isolation, then centrifuged at 3000 rpm for 20 minutes of blood draw, and finally placed on ice. Serum samples were measured at 1:4 dilution. NfL concentrations in serum were measured using a commercial ELISA (NF-light® ELISA, Uman Diagnostics, Umeå, Sweden). We used Anti NF-L monoclonal antibody (mAB) as a capture antibody and a biotin-labeled Anti NF-L mAB as the detection antibody. All samples measured blinded. ELISA readings were converted to units per milliliter by using a standard curve constructed by calibrators (Bovine lyophilized NfL obtained from UmanDiagnostics).

### 2.5 Accuracy, speed, and ICA summary score calculations

Participants’ responses to each image and their reaction times (i.e. time between image onset and response) are recorded and used to calculate their overall accuracy and speed. Speed and accuracy are then used to calculate an overall summary score, we refer to as the ICA score.

***Accuracy*** is simply defined as the number of correct categorisations divided by the total number of images, multiplied by a 100.

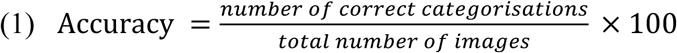

***Speed*** is defined based on participant’s response reaction times in trials they responded correctly.

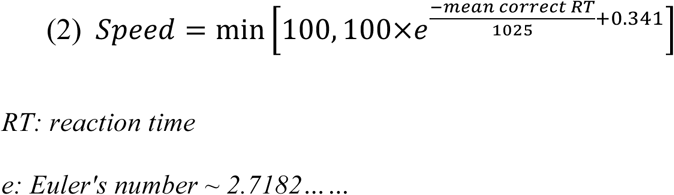

Speed is inversely related with participants’ reaction times; the higher the speed, the lower the reaction time.

#### Preprocessing

We used boxplot to remove outlier reaction times, before computing the ICA score. Boxplot is a non-parametric method for describing groups of numerical data through their quartiles; and allows for detection of outliers in the data. Following the boxplot approach, reaction times greater than q3 + w * (q3 - q1) or less than q1 - w * (q3 - q1) are considered outliers. q1 is the lower quartile, and q3 is the upper quartile of the reaction times. Where “w” is a ‘whisker’; w = 1.5. The number of reaction-time data-points removed by the boxplot can vary case by case, but in all participants it was less than 40% of their data (data refer to the number of images they observed).

The ***ICA summary score*** is a combination of accuracy and speed, defined as follows:

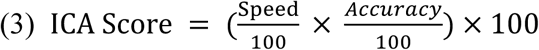

### 2.6 ICA’s artificial intelligence (AI) engine

ICA’s AI engine used in this study was a multinomial logistic regression (MLR) classifier trained based on the set of ICA features extracted from the ICA test for each participant. These features included, the ICA score, and the trend of speed and accuracy during the test (i.e. whether the speed and/or accuracy were increasing or decreasing during the time-course of the test). The classifier also took subject’s age, gender and education in order to match subjects with similar demographics.

**Multinomial logistic regression classifier (MLR)**^34^ is a supervised regression-based learning algorithm. The learning algorithm’s task is to learn a set of weights for a regression model that maps participants’ ICA test output to classification labels.

The basic difference between ICA’s classification of patients (using the AI engine) and the conventional way of defining an optimal cut-off value for classification is the dimensionality (or the number of features) used to make the classification. For example, in a conventional assessment tool, an optimal cut-off value is defined based on the test score. This is a one-dimensional classification problem, and there is only one free parameter to optimize, therefore less flexibility to learn from more data. In ICA, however, the test returns a rich set of features (we have one reaction-time and accuracy per each image). ICA score is the most informative summary score, but on top of this, we used a classifier to find the optimum classification boundary in the higher dimensional space. There are more free parameters here to optimize and therefore, the classifier can benefit from more data to best set these parameters for achieving a higher accuracy.

## 3. Results

### 3.1 Convergent validity with BICAMS, and sensitivity to MS

In substudy 1, we assessed convergent validity by examining the correlation between scores on the ICA test and the BICAMS battery (i.e. SDMT, BVMT-R and CVLT-II). Figure 2 presents scatterplots examining the relationship between BICAMS and ICA test performance. A high level of convergent validity is demonstrated between ICA and BICAMS. Within the BICAMS battery, SDMT had the highest correlation with the ICA test for HC (Pearson’s r = 0.81), MS (r = 0.71), and combined (r = 0.82) groups. Scatterplots show ICA vs. BICAMS correlation separately for MS and HC; combining results from both groups (n = 174 total), we find a correlation of 0.82 with SDMT, 0.71 with CVLT-II, and 0.60 with BVMT-R. Correlations between ICA’s speed and accuracy components with the BICAMS battery are also reported in Table 3.

**Figure 2.**
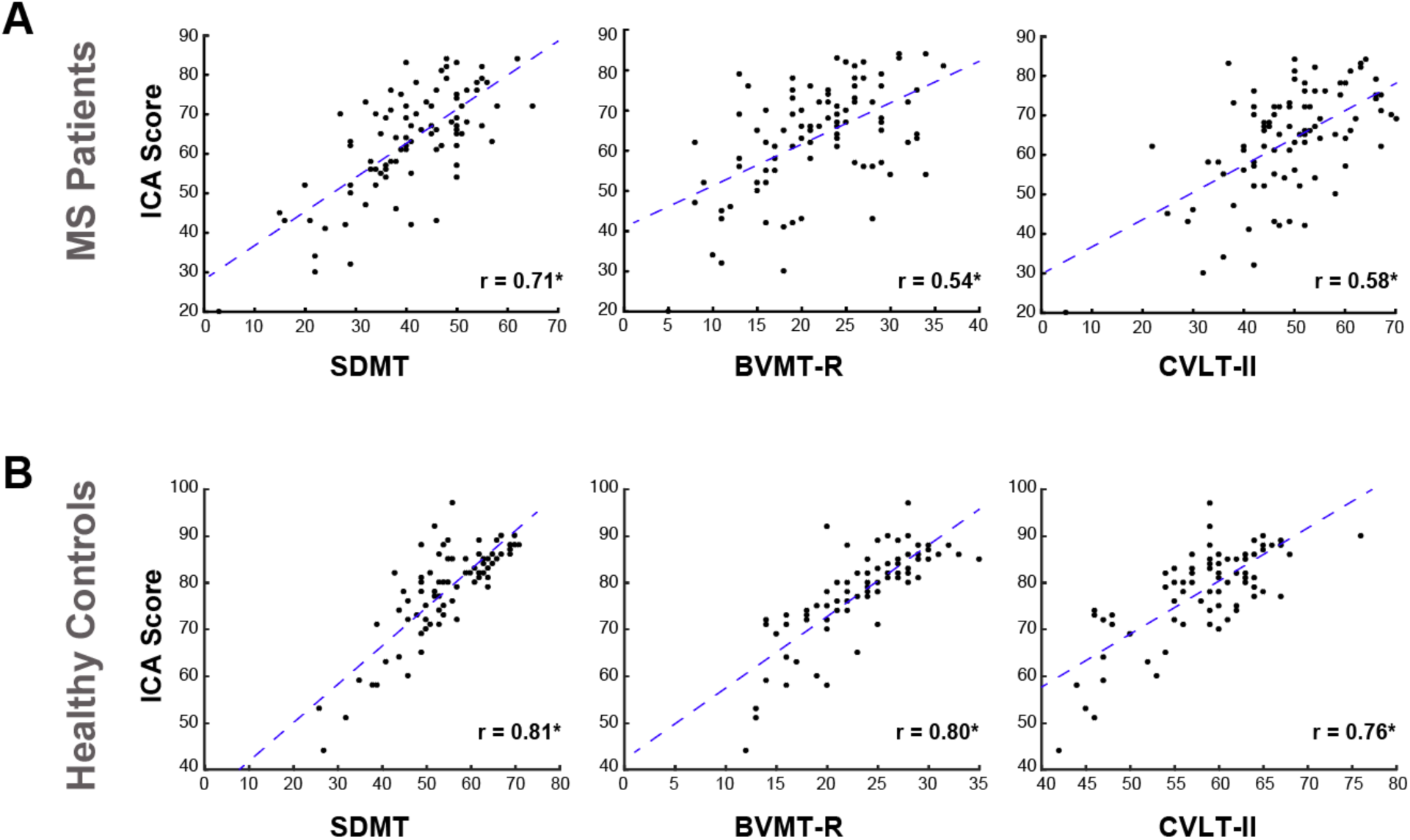
Correlation between BICAMS and ICA for (A) MS patients and (B) healthy controls. Each scatter plot shows the ICA score (y axis) vs. one of the tests in BICAMS (x axis). Each blue dot indicates an individual; the blue dashed lines are results of linear regression, fitting a linear line to the data in each plot. For each plot, the Pearson correlation between ICA and a BICAMS test is written on the bottom-right. If we combine the data from MS patients and healthy controls (n = 174 total), the ICA vs. BICAMS correlations will be the following: correlation with SDMT: 0.82; BVMT-R: 0.60; CVLT-II: 0.71. ICA: Integrated Cognitive Assessment; SDMT: Symbol Digit Modalities Test; BVMTR: Brief Visual Memory Test–Revised; CVLT-II: California Verbal Learning Test -2nd edition. Stars (*) show significant correlation at p<10^−8^.

**Table 3.** Speed and accuracy correlations with BICAMS.

|  | SDMT | BVMT-R | CVLT-II |
| --- | --- | --- | --- |
| Speed | $r=0.66 *$ | $r=0.42 *$ | $r=0.52 *$ |
| Accuracy | $r=0.55 *$ | $r=0.46 *$ | $r=0.52 *$ |

Pearson correlations between the BICAMS battery and speed, accuracy components of the ICA test across all participants. (\* shows statistical significance at $p < 10^{-6}$ )

To compare sensitivity of BICAMS and ICA in detecting MS dysfunctions, we compared mean test scores in MS and HC groups separately for BICAMS battery of tests and the ICA test (Table 3). Within the BICAMS battery, SDMT and CVLT-II could differentiate between HC and MS patients (Table 3). The scores on both SDMT and CVLT-II were significantly lower for the MS patients compared to the HC group. However, there was no significant difference between BVMT-R scores of the HC and MS groups. These results are consistent with previous findings showing that SDMT has a better sensitivity in detecting MS compared to other tests within the BICAMS battery ^5,35^.

### 3.2 ICA accuracy in detecting cognitive impairment

Participants were categorized into cognitively intact and cognitively impaired by the neurologist in their outpatient visit, based on their presenting complaint and the BICAMS cognitive assessment. Following this procedure 45% of MS patients were classified with cognitive impairment. Using an ROC curve (Figure 3), we then assessed the accuracy of the ICA’s AI engine (i.e. MLR classifier) in discriminating cognitively healthy from cognitively impaired individuals (Figure 3, area under curve (AUC) = 95.1 %, sensitivity = 82.9 %, and Specificity = 96.1%.)

**Figure 3.**
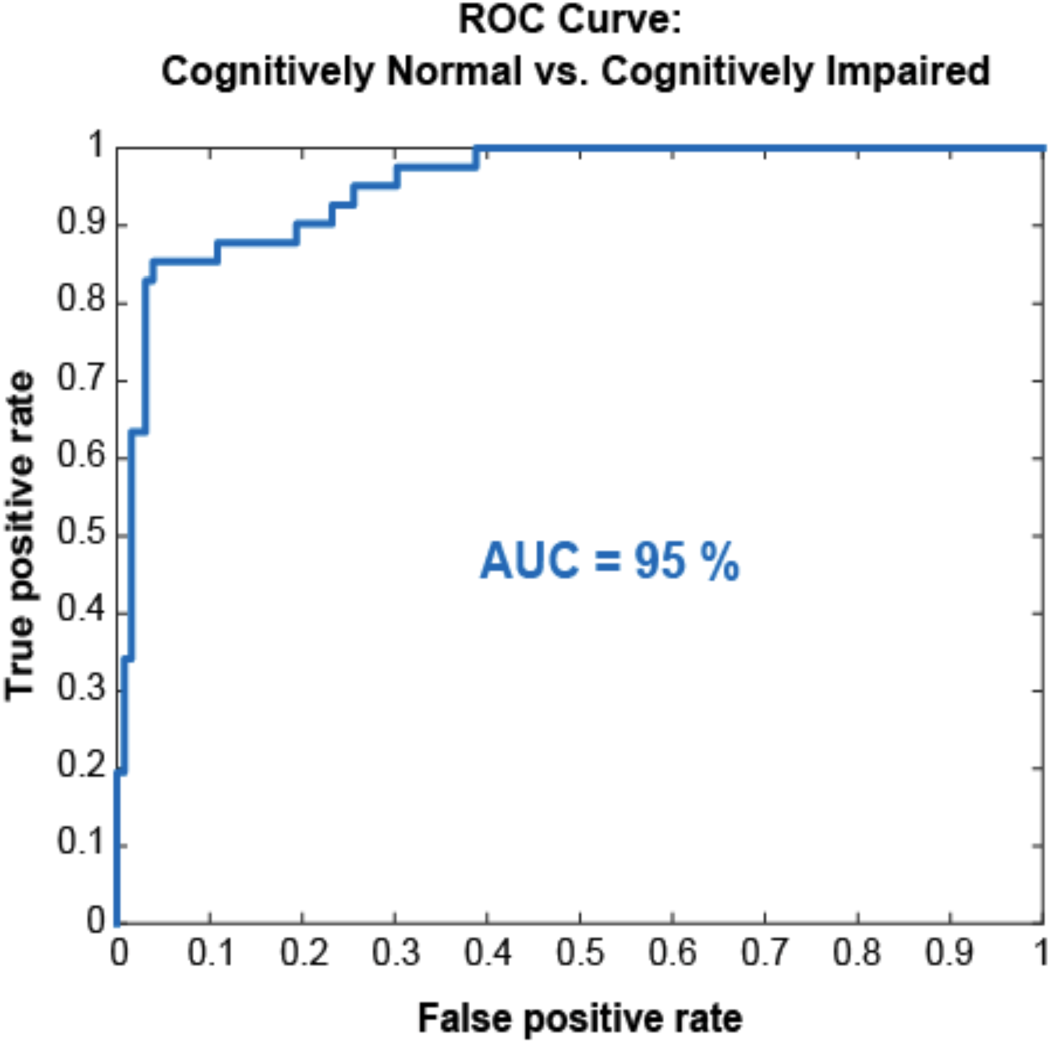
ROC curve for the ICA test in discriminating cognitively impaired from cognitively healthy individuals. A multinomial logistic regression classifier was trained based on the ICA test output, and tested using leave-one-out cross-validation. AUC =95.1%; Sensitivity = 82.9 %; Specificity = 96.1%.

### 3.3 ICA and SDMT correlations with Neurofilament light (NfL)

Neurofilament light (NfL) is a promising fluid biomarker of disease progression for various brain disorders, such as Alzheimer’s Disease and Multiple Sclerosis ^36,37^. In substudy 2, we demonstrated that there is a strong correlation between ICA score and the level of serum NfL (Figure 4A). For comparison, on the same set of MS participants, SDMT correlations with NfL is also reported (Figure 4B). SDMT and ICA were both administered in the same session.

**Figure 4.**
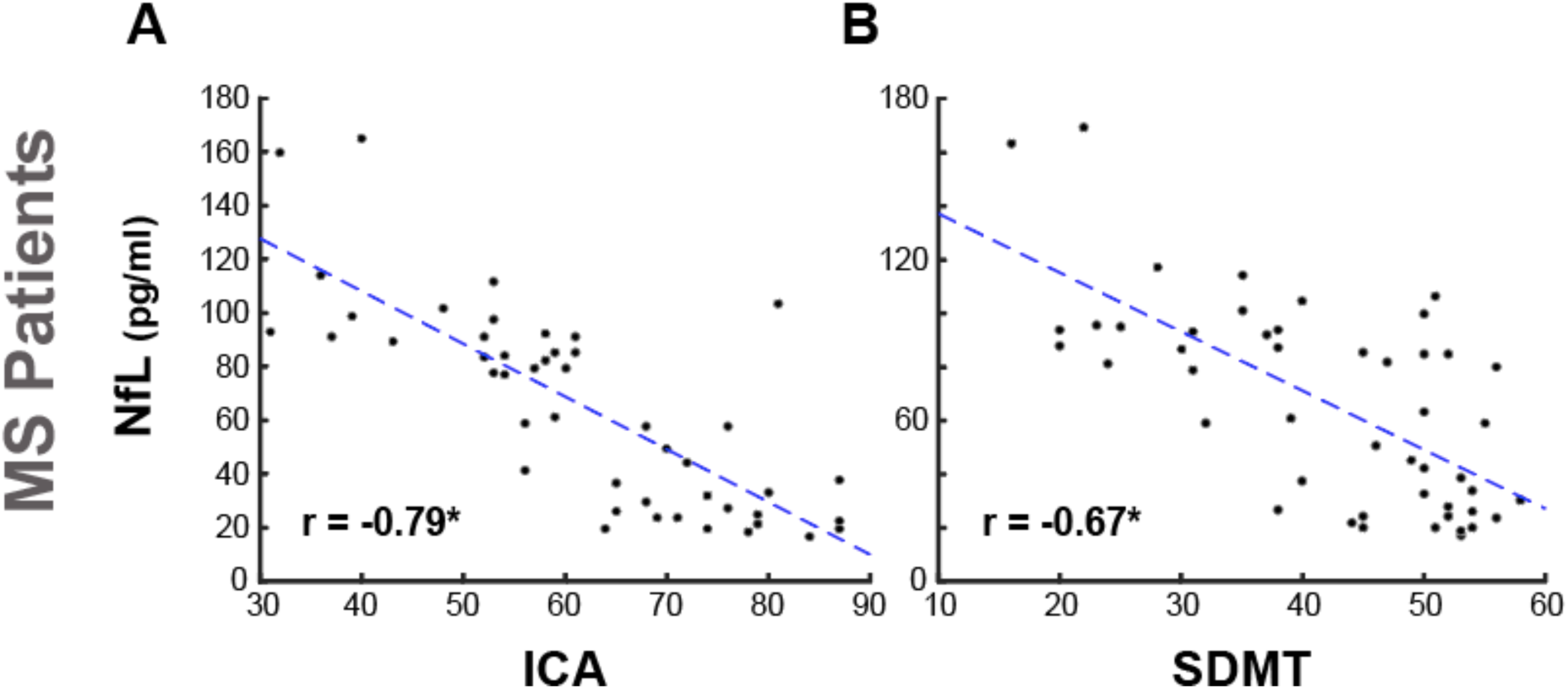
ICA correlation with severity of neural damage, as measured by serum NfL. Each scatter plot shows the NfL level in serum (y axis) vs. ICA or SDMT (x axis). Each blue dot indicates an individual; the blue dashed lines are results of linear regression, fitting a linear line to the data in each plot. For each plot, the Pearson correlation between NfL level and the reference cognitive test is written on the bottom-left. Stars (*) show significant correlations at p<10^−6^.

### 3.4 ICA test-retest reliability and absence of a learning bias

Test-retest reliability was measured by computing the Pearson correlation between the two ICA scores. R values for test-retest correlation are considered adequate if >0.70 and good if >0.80 ^38^. Figure 5 presents scatterplots of ICA performance comparing 1st administration versus 2nd administration of the test for the HC, MS, and combined groups. Test–retest reliability was high, with correlation values in the range between 0.91 and 0.94.

**Figure 5.**
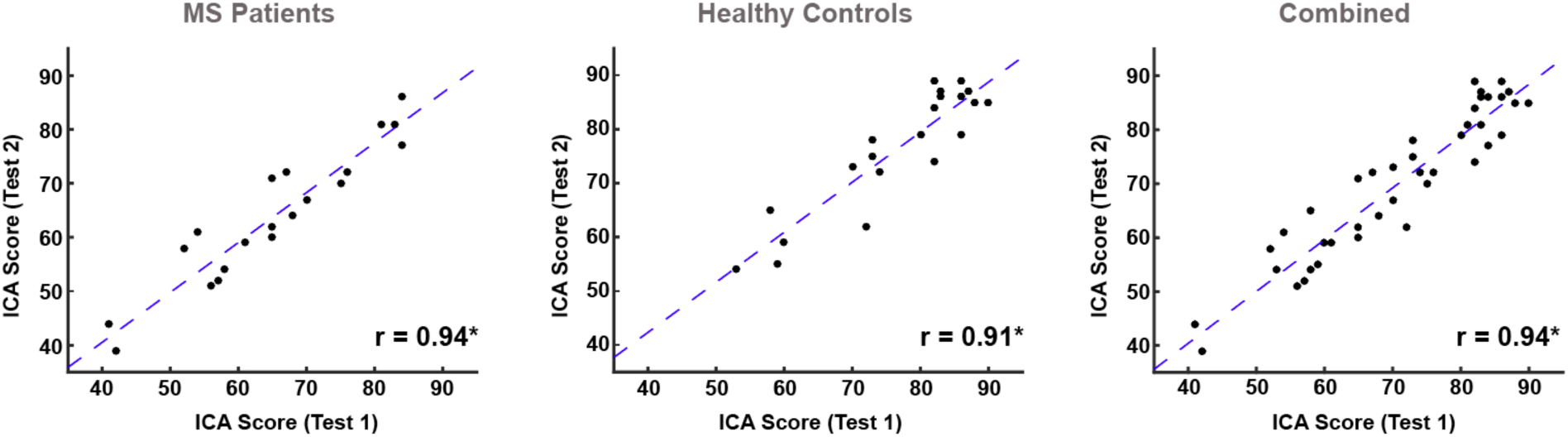
Test-retest reliability scatter plots for the ICA test. Scatterplots are presented comparing ICA scores at Time 1 versus Time 2 administrations for the MS, HC, and combined groups. The gap between the 1^st^ and the 2^nd^ administration of the ICA test was 5 weeks (+-15 days). Reliability is calculated using Pearson’s r. The test-retest reliability for the SDMT test was: r (combined) = 0.97; r (HC) = 0.98; r (MS) = 0.97. Stars (*) indicate statistical significance at p < 10^−8^.

In the subgroup of participants (21 MS, and 22 HC) who took the ICA and SDMT for a second time, we studied whether they could systematically get a better performance due to a previous exposure to either of the tests. This is called a learning bias (also referred to as practice effect). As shown in Table 5, comparing the first and second administration of the ICA and SDMT tests, ICA showed no learning bias. However, we see an improvement in participant’s average SDMT score. This improvement in SDMT score (i.e. learning bias) was statistically significant in the HC group, but not in the MS group.

**Table 4.** Mean ICA and BICAMS scores per group.

|  | MS (n=91) |  | HC (n=83) |  |  |  |  |
| --- | --- | --- | --- | --- | --- | --- | --- |
| BICAMS | mean | SD | mean | SD | Difference | Cohen's d | p-value |
| SDMT | 40.04 | 11.02 | 54.73 | 9.77 | 14.69 | 1.31 | <10<sup>-14</sup> |
| BVMT-R | 21.89 | 6.95 | 23.69 | 5.17 | 1.80 | 0.29 | =0.0565 |
| CVLT-II | 48.96 | 11.13 | 58.28 | 6.59 | 9.32 | 1.00 | <10<sup>-9</sup> |
| ICA |  |  |  |  |  |  |  |
| ICA score | 63.67 | 13.30 | 78.43 | 9.86 | 17.76 | 1.26 | <10<sup>-13</sup> |
| Accuracy | 84.97 | 11.66 | 89.57 | 5.79 | 4.60 | 0.50 | =0.0014 |
| Speed | 74.54 | 12.26 | 87.76 | 10.11 | 13.22 | 1.16 | <10<sup>-12</sup> |

Mean and standard deviations (SD) for BICAMS test scores and the ICA test scores are compared for MS patients versus healthy controls (HC). The ICA score is a composite score made of both speed and accuracy of participants in ICA's rapid visual categorization task. P-values come from a two-sample t-test.

**Table 5.** Learning bias (practice effect) for ICA and SDMT.

| Group | Test Name | Test 1<br>[mean ± SD] | Test 2<br>[mean ± SD] | Paired t-test | | |
| --- | --- | --- | --- | --- | --- | --- |
|  |  |  |  | difference | Cohen's d | p-value |
| MS (*n*=21) | ICA | 64.76 ±12.7 | 64.66 ±12.2 | -0.09 | -0.02 | 0.91 |
| | SDMT | 43.6 ±10 | 44.2 ±11 | 0.66 | 0.25 | 0.25 |
| HC (*n*=22) | ICA | 77.04 ±11 | 76.86 ±11 | -0.18 | -0.07 | 0.73 |
| | SDMT | 55.13 ±12 | 56.36 ±12.3 | 1.22 | 0.50 | 0.02* |

Mean and standard deviations (SD) of SDMT and ICA scores are compared between the two administrations of the tests for each group of participants. Only SDMT in healthy subjects showed a significant practice effect.

### 3.5 ICA correlation with EDSS, age and education

To further characterize the ICA score and its relationship with other measures from the MS patients, we calculated the correlation between ICA score and patients’ EDSS, age and education (Table 6). Both BICAMS and ICA scores were negatively correlated with patients’ EDSS, demonstrating an inverse relation between disability scale and cognitive performance. For all the tests, we also observed a decrease in performance as the age increases, showing the effect of aging on cognitive performance. All tests were correlated with participant’s level of education, with ICA having the lowest correlation.

**Table 6:** Age/EDSS/Education vs. BICAMS/ICA.

|  |  | **BICAMS** |  |  |  |
| --- | --- | --- | --- | --- | --- |
|  |  | **SDMT** | **BVMT-R** | **CVLT-II** | **ICA** |
| **Correlation with** | **EDSS** | -0.41<br>($p < 10^{-4}$) | -0.26<br>($p < 0.05$) | -0.33<br>($p < 0.001$) | -0.58 ($p < 10^{-8}$) |
| | **Education** | 0.50<br>($p < 10^{-6}$) | 0.34<br>($p < 0.001$) | 0.31<br>($p < 0.01$) | 0.25 ($p < 0.05$) |
| | **Age** | -0.38<br>($p < 10^{-4}$) | -0.50<br>($p < 10^{-5}$) | -0.42<br>($p < 10^{-4}$) | -0.49 ($p < 10^{-6}$) |

The table shows Pearson Correlations of the ICA score and the BICAMS battery of tests with MS patients' EDSS score, education in years, and their age. EDSS: Expanded Disability Status Scale; BICAMS: Brief International Cognitive Assessment for MS; SDMT: Symbol Digit Modalities Test; BVMT-R: Brief Visual Memory Test-Revised; CVLT-II: California Verbal Learning Test -2nd edition; ICA: Integrated Cognitive Assessment.

## 4. Discussion

In this validation study, we demonstrate that the ICA test has convergent validity with BICAMS, with an excellent test-retest reliability comparable to that reported for SDMT ^9^. ICA is a visuo-motor test and primarily tests information processing speed (IPS) and engages higher visual areas in the brain for semantic processing (i.e. animal vs. non-animal). Comparing speed versus accuracy in the ICA test (Table 4), speed seems to play a more significant role in discriminating MS patients from HC participants. This corroborates findings from other studies suggesting slower speed of information processing as a key deficit in multiple sclerosis ^39^. IPS impairment underlies other areas of cognitive dysfunction ^14,40^. This is because the speed with which an individual performs a cognitive task is not simply an isolated function of the processes required in that task, but also a reflection of their ability to rapidly carry out many different types of processing operations. In the case of ICA, these operations include transferring visual information through retina to higher level visual areas (i.e. sensory speed), processing the image representation in the visual system to categorize it into animal or non-animal (i.e. cognitive speed), and then translating this into a motor response (i.e. motor speed).

In contrast to currently standard cognitive tests, whereby stimuli are language-dependent, the presented stimuli in the ICA test are natural images that contain universally recognizable images of animals or objects, thus making the test intrinsically language-independent. Furthermore, participants’ responses only involve tapping on the left or right side of an iPad, making it totally independent of participants’ knowledge of Arabic numerals or any other symbolic representation of numbers (as used in SDMT) or alphabets and words (as in CVLT-II), or the drawing ability of a participant when drawing shapes (as in BVMT-R). This makes the ICA test more suitable for wider international use, and less dependent on lingual, educational, and demographic differences.

Comparing ICA with other reliable iPad-based cognitive tests, such as the electronic implementations of SDMT ^41,42^, we would like to highlight two main differences: a) The ICA test takes advantage of an AI platform, thus the capacity to learn from big data and further fine tune its multidimensional classification boundaries when it comes to see new training cases. b) ICA did not show a learning bias in this study and a previous study ^18^, as opposed to the learning bias reported for the iPad-based SDMT (i.e. PST) ^41^. This is because, in ICA, the images are shown in random order and they are presented only for 100 ms, making it very difficult for participants to learn the test.

For making an early diagnosis of MS and monitoring the disease progression, we need reliable biomarkers. NfL has been shown to be a promising fluid biomarker of disease progression for various brain disorders, including MS ^10,43^. Increased levels of NfL are shown to correlate with the severity of neural damage ^44^. In substudy 2, we demonstrated a strong association between ICA score and NfL in MS patients. This is particularly of interest given the totally non-invasive nature of the ICA test; and suggests the use of ICA as a digital biomarker of cognition in MS. For more frequent cognitive assessments, digital biomarkers have an advantage over fluid biomarkers, given their lower cost, accessibility, the possibility of remote administration and easier integration into routine clinical practice.

Some limitations encountered in this study include the lack of NfL data from the healthy control group, and the absence of neuroimaging markers of disease activity. Future studies are needed to investigate the link between ICA test results and neuroimaging biomarkers of MS, in particular, the correlation between T2 lesion load and ICA score would be of interest.

Our results provide evidence for using ICA in neurology outpatient clinics as an accurate tool for assessing cognitive impairment in MS. Digital biomarkers of cognition (such as ICA) can be easily used to measure changes in cognitive performance relative to a baseline, which subsequently paves the way for using cognition as a marker of disease progression and treatment efficacy in MS. Given the absence of a learning bias in ICA, the test can be considered suitable for frequent monitoring of cognitive performance, allowing clinicians to measure cognitive decline, as well as potential treatment efficacy. The test further has the potential to be used for remote (i.e. home-based) monitoring of cognitive performance. Future longitudinal studies need to test this on large populations.

## Funding

SMKR was funded by a return home fellowship grant from the Iranian National Elite Foundation. Cognetivity ltd covered the costs for purchasing BICAMS. Other costs were covered by the Royan Institute internal funds to the investigators SMN, and SMKR.

## Authors Contributions

SMN and SMKR and CK conceived and designed the study. SMN, MS, and MK did the data collection and patient recruitment. The data were analyzed by MS and MK, under the supervision of MSN and SMKR. All authors were involved in writing the manuscript.

## Declaration of Conflicting Interests

SMKR serves as the chief science officer at Cognetivity ltd; CK serves as Chief Medical Officer at Cognetivity Ltd. Other authors declared no potential conflicts of interest.

